# Chromenone derivatives as novel pharmacological chaperones for retinitis pigmentosa-linked rod opsin mutants

**DOI:** 10.1101/2022.04.05.487228

**Authors:** Joseph T. Ortega, Andrew G. McKee, Francis J. Roushar, Wesley D. Penn, Jonathan P. Schlebach, Beata Jastrzebska

**Affiliations:** Department of Pharmacology and Cleveland Center for Membrane and Structural Biology, School of Medicine, Case Western Reserve University, 10900 Euclid Ave., Cleveland, OH 44106, USA; Department of Chemistry, Indiana University, 800 E. Kirkwood Ave., Bloomington, IN 47405-7102, USA

**Author notes:** **CORRESPONDENCE**, Beata Jastrzebska, Ph.D., Department of Pharmacology, School of Medicine, Case Western Reserve University, 10900 Euclid Ave., Cleveland, OH 44106-4965, USA, Jonathan P. Schlebach, Ph.D., Department of Chemistry, Indiana University, 800 E. Kirkwood Ave., Bloomington, IN 47405-7102, USA.

**Keywords:** chaperone, mutant, photoreceptor, retinal degeneration, retinitis pigmentosa, rhodopsin

## Abstract

The correct expression of folded, functional rhodopsin (Rho) is critical for visual perception. However, this seven-transmembrane helical G protein-coupled receptor (GPCR) is prone to mutations with pathological consequences of retinal degeneration in retinitis pigmentosa (RP) due to Rho misfolding. Pharmacological chaperones that stabilize the inherited Rho variants by assisting their folding and membrane targeting could slow the progression of RP. In this study, we employed virtual screening of synthetic compounds with natural product scaffold in conjunction with *in vitro* and *in vivo* evaluations to discover a novel chromenone-containing small molecule with favorable pharmacological properties that stabilizes rod opsin. This compound reversibly binds to unliganded bovine rod opsin with an EC_50_ value comparable to the 9-*cis*-retinal chromophore analog and partially rescued membrane trafficking of multiple RP-related rod opsin variants *in vitro*. Importantly, this novel ligand of rod opsin was effective *in vivo* in murine models, protecting photoreceptors from deterioration caused either by bright light or genetic insult. Together, our current study suggests potential broad therapeutic implications of the new chromenone-containing non-retinoid small molecule against retinal diseases associated with photoreceptor degeneration.

## INTRODUCTION

Inherited retinal degeneration associated with mutations in the rhodopsin (*RHO*) gene is a common cause of a spectrum of blinding diseases called retinitis pigmentosa (RP), for which therapy is currently not available (1,2). Many of the ~150 identified mutations are associated with impaired trafficking of the mutated Rho protein due to their instability, misfolding, and aberrant binding of the native chromophore 11-*cis*-retinal (2,3). Expression of defective Rho results in the formation of disorganized rod outer segment discs, which leads to rod photoreceptor degeneration followed by cone photoreceptors death (1). In the human population, the substitution of Pro23 to His in the *RHO* gene occurs the most frequently and accounts for about one-third of RP cases (1,4). The folding and membrane targeting of this mutant could be stabilized by the native chromophore and its analog 9-*cis*-retinal *in vitro* (5–7). The beneficial effect of treatment with high doses of vitamin A, a precursor of 11-*cis*-retinal, was demonstrated in patients with RP (8,9). However, limited efficacy, chemical instability, and toxicity of the photo-metabolites limit the pharmacological potential of retinal compounds (10–12). The newly synthesized receptors exist in the photoreceptor cells in the ligand-free state perhaps until they reach the rod outers segments (ROS). Thus, genetically impaired rod opsin is specifically vulnerable within the biosynthetic pathway, where the supply of 11-*cis*-retinal is limited. Therefore, the development of non-retinoid compounds that could support opsin stability and folding during biosynthesis is highly attractive (10,11,13).

Rhodopsin’s chromophore-binding pocket is flexible and large enough to accommodate other molecules than its native 11-*cis*-retinal ligand (14). Thus, targeting the opsin’s orthosteric site could allow for the identification of novel non-retinoid small molecules that stabilize rod opsin and stimulate trafficking of misfolding mutants from the endoplasmic reticulum (ER) to the rod outer segments. Indeed, several such non-retinoid stabilizers with pharmacological chaperone potential were identified recently through virtual and high-throughput screenings of small molecule libraries (15–17). Alternatively, we found that compounds with natural product scaffold could accommodate within the opsin’s orthosteric site, providing an option to screen for new chemical correctors of pathogenic variants (18). These biologically active polyphenolic compounds, specifically flavonoids, could enhance the stability of unliganded rod opsin, and improve the membrane expression and function of P23H mutant *in vitro* (18,19). Moreover, as we discovered treatment with quercetin could prevent retinal degeneration in mouse models of acute light damage and importantly, slowed down the progression of photoreceptors death in the P23H Rho *knock-in* mice, a model of autosomal dominant RP (adRP) (19,20).

In this current follow-up study, we screened a natural product-derived synthetic molecule library aiming to discover novel opsin ligands with chaperone-like activities towards pathogenic mutants. This library included oxygen heterocycles, polyketides, flavonoids, terpenoids, alkaloids, steroids, and other derivatives, providing additional chemical diversity with higher synthesis viability for future lead optimization. The *in silico* analysis conducted in this study led to the identification of five new compounds with a common chemical scaffold containing a chromenone motif. Chromenones have become an important class of oxygenated heterocycles due to their broad spectrum of biological activities (21–24). Interestingly, several chromenone-containing compounds were identified as selective potent agonists of the orphan G protein-coupled receptor 35 (GPCR35) and GPCR55 (25–27). The major advantage of the chromenone moiety is its accessibility for the compound chemical modification to improve its biological effects (28).

Our results show that some of these compounds could bind to unliganded rod opsin and modulate the function of this receptor. Moreover, these compounds showed a stabilizing effect for the pathogenic P23H rod opsin during biosynthesis resulting in the improved transport of this variant to the cell surface. Furthermore, we surveyed the potential of these new chromenone derivatives to rescue membrane targeting of other misfolding Rho variants by using a recently described deep mutational scanning approach, identifying a panel of mutants most amenable to be rescued by one of these molecules (29–31).

To validate the therapeutic potential of the identified lead compound, we evaluated its activity in two mouse models of retinal degeneration, *Abca4^-/-^Rdh8^-/-^* mice and the P23H Rho *knock-in* mice. In *Abca4^-/-^Rdh8^-/-^* mice exposure to acute light triggers photoreceptors death caused by the excessive local concentrations of unliganded opsin and toxicity of retinal byproducts (32,33). On the other hand, the P23H Rho *knock-in* mice resemble many features of human RP (34). Both mouse models were proven to be excellent models to evaluate drug-like activity of retinal and non-retinal opsin ligands against retina degeneration (16,19,20,35). The identified here chromenone-containing compound showed beneficial effects for retina health in both *Abca4^-/-^ Rdh8^-/-^* and P23H Rho mice, suggesting its potential as a therapeutic lead compound with broad application for retinal diseases related to photoreceptor degeneration.

## MATERIAL AND METHODS

### Chemicals and Reagents

Peanut agglutinin (PNA) and Alexa Fluor 488-conjugated streptavidin were purchased from Vector. 4969-Diamidino-2-phenyl-indole (DAPI) for the nuclear staining was purchased from Life Technologies (Grand Island, NY). Dimethylsulfoxide (DMSO) was obtained from Sigma (St. Louis, MO). EDTA-free protease inhibitor cocktail tablets were purchased from Roche (Basel, Switzerland). 9-*cis*-retinal was purchased from Sigma. Polyvinylidene difluoride (PVDF) membrane was obtained from Millipore (Burlington, MA). Compounds CR1-CR5 were purchased from OTAVAchemicals Ltd. (Ontario, Canada).

### *In silico* Compound Screening

The coordinates of the ligand-free bovine opsin at 2.9 Å resolution were obtained from PDB ID: 3CAP. One monomeric unit of the protein was selected. Then, co-crystallized molecules and all crystallographic water molecules were removed from the coordinate set. Hydrogen atoms were added, and partial charges were assigned to all atoms. The protein was then submitted to the restrained molecular mechanics refinement with NAMD 2.12 software, using the CHARMM22 force field. The coordinates for the ligand-binding pocket located in the orthosteric site were selected based on the results obtained for quercetin as reported in (18). The screening library was built using the 3D structures of the compounds from the OTAVA database. The library was evaluated using molecular docking carried out with VINA/Vega 3.1.0.21 software (36). The results were prioritized according to the predicted binding free energy in kcal/mol, and the best 100 compounds were then re-docked. The results obtained from docking were visualized with the Biovia Discovery Studio Visualizer 17.2.0 software. These results were suitable for an evaluation to assess the position into the orthosteric site, interactions formed between the compound and residues in the pocket and coverage of different chemical scaffolds to avoid overflow the library with similar chemical compounds. Quercetin was used as a positive control for the run and to define the cutoff for binding free energy to select the hit compounds. The top 25 compounds with lower binding free energy than quercetin were selected. The docking results obtained were then rescored by running ten times each compound. Then, the binding poses and interactions (location into the binding site; number and type of interactions formed; different chemical scaffolds) were analyzed by visual inspection process, using Biovia Discovery Studio Visualizer 17.2.0 software.

To confirm the stability of the interaction between the compound and opsin, the opsin-compound complexes obtained from molecular docking were used to perform the molecular dynamic simulations using VegaZZ software. Protein-compound complexes were inserted in a membrane of POPC (1-palmitoyl-2-oleoyl-sn-glycerol-3-phosphocholine) and immersed in a box (10×10×10 Å) filled with water molecules. Counter ions were added to neutralize charges. The complex energies were minimized as described above and NPT simulations were carried out using NAMD and CHARMM force field. The length of each simulation was 100 ns. The obtained results were expressed as root-square mean deviation (RMSD).

### Analysis of Pharmacokinetic Drug Properties

A comprehensive analysis of physicochemical descriptors, parameters related to administration, distribution, metabolism, and elimination (ADME), drug-like nature of the identified compounds was carried out using SWISSADME tools (37). These tools were accessed through the website at http://www.swissadme.ch.

### Preparation of Opsin Membranes

Rod outer segment (ROS) membranes were isolated from frozen bovine retinas under dim red light according to the protocol described previously (38). The membrane-associated proteins were washed out by four washes with a hypotonic buffer composed of 5 mM HEPES, pH 7.5, and 1 mM EDTA. Membranes were gently homogenized and pelleted by centrifugation at 25,000xg for 25 min. The final membrane pellet was suspended in 10 mM sodium phosphate, pH 7.0, and 50 mM hydroxylamine at Rho concentrations of ~2 mg/ml. The membranes then were exposed to light with a 150 Watt bulb for 30 min at 0 °C. Next, these membranes were centrifuged at 16,000 x g for 10 min. The supernatant was discarded and the membrane pellet was washed twice with 10 mM sodium phosphate, pH 7.0, containing 2% bovine serum albumin (BSA) followed by 4 washes with 10 mM sodium phosphate, pH 7.0, and 2 washes with 20 mM bis-tris-propane (BTP), pH 7.5, and 100 mM NaCl. After each wash, the membranes were pelleted by centrifugation at 16,000xg for 10 min at 4 °C.

### Compounds-Opsin Binding and Pigment Regeneration

Washed ROS membranes were resuspended in the buffer consisting of 20 mM BTP, pH 7.5, and 100 mM NaCl at 5 μM final concentration of opsin. The compounds were added to these membranes at 10 μM concentration and incubated for 30 min at a rotating platform at RT followed by the addition of 5 μM 9-*cis*-retinal and incubation for 15 min. Alternatively, membranes were incubated only with 9-*cis*-retinal. Then, membranes were solubilized with 20 mM dodecyl-β-D-maltopyranoside (DDM) for 5 min at RT, followed by centrifugation at 16,000xg for 5 min at 4 °C. The UV-visible spectra were measured immediately with a Cary 60 spectrophotometer.

Alternatively, compounds at 1, 10, and 100 μM concentrations were incubated with ROS membranes followed by membrane solubilization with 20 mM DDM for 5 min at RT. Solubilized opsin was cleared by centrifugation at 16,000xg for 5 min at 4 °C. Then, 5 μM 9-*cis*-retinal was added to the sample and UV-visible spectra were measured every 2 min for 60 min at 20 °C. Each condition was repeated three times. The absorbance at 485 nm was plotted as a function of time and a time course of pigment regeneration was fitted to a second-order exponential decay to calculate the rates of regeneration and the apparent half-lives of isorhodopsin (isoRho) regeneration.

### UV-visible Spectroscopy of Opsin, Rho and isoRho

The concentration of Rho and opsin measured within ROS membranes was determined after membrane solubilization with 20 mM DDM and pelleting insoluble material by centrifugation at 16,000xg for 15 min at 4 °C using a UV-visible spectrophotometer (Cary 60, Varian, Palo Alto, CA). The absorption coefficients ε_500nm_=40,600 M^-1^cm^-1^ was used for Rho, ε_280nm_=81,200 M^-1^cm^-1^ for opsin, and ε_485nm_=43,600 M^-1^cm^-1^ for isoRho concentration calculations (39,40).

### Fluorescence Spectroscopy

The quenching of the intrinsic Trp fluorescence was also used to confirm the binding of the new compounds into the opsin’s orthosteric binding pocket (16,18). Trp opsin’s fluorescence was monitored before and after addition of the new compound at concentrations of 0.1, 0.2, 0.3, 0.4, 0.5, 1.0, and 1.5 μM. The emission spectra were recorded with an FL 6500 Fluorescence Spectrometer at 20 °C between 300 and 420 nm after excitation at 295 nm. The excitation and emission slit bands were set at 5 and 10 nm, respectively. Changes in the intrinsic Trp fluorescence at 330 nm (Δ*F*/*F*_0_, where *ΔF* is the difference between the initial Trp fluorescence (*F*_0_) and fluorescence recorded upon addition of the compound) were plotted as a function of the ligand concentration. The ligand-binding curves were fitted and half-maximal effective binding concentrations (EC_50_) were calculated using the GraphPad Prism 7.0 software. All the measurements were performed in triplicate. All experimental data were corrected for the samples background and self-absorption at excitation and emission wavelengths (inner filter effect correction). Each condition was performed in triplicate and the experiment was repeated.

### Thermal Shift Assay

Twenty μl of opsin membranes suspended in 20 mM BTP, pH 7.5, and 100 mM NaCl at a concentration of 0.01 mg/ml were loaded into a 96-well plate (Applied Biosystem). Compound were then added at concentrations of 0.1, 1, 10, 100 nM and 1 and 10 μM to each well and incubated for 1 h at 4 °C. Next, five μl of the BiFC probe from 10 mM stock solution in DMSO was added to each well. Opsin membranes without treatment were used as a control. The plate was sealed with a ClearSeal film (HR4-521, Hampton Research) and incubated for 10 min on ice prior to measuring changes in the sample fluorescence. All measurements were performed with a StepOnePlus Real-Time PCR System (Applied Biosystems). The melting curve experiments were recorded using the StepOne software version 2.3. The fluorescence in the SYBR, FAM, and ROX channels was recorded for each sample. The run was set to cool the plate to 4 °C within 10 s, kept at 4 °C for 1 min, and then increased 1 °C per min in a step-and-hold manner up to 99.9 °C. The multicomponent data were exported to a Microsoft Excel sheet and analyzed with the Prism 6.0 software. The melting temperatures (Tm) of bovine Rho and opsin within the membranes were 71.9 and 54.8 °C, respectively (18,41). Each condition was performed in triplicate and the experiment was repeated.

### G_t_ Activation Assay

G_t_ was extracted and purified from ROS membranes isolated from a hundred dark-adapted bovine retinas according to the protocol described in (18,42). The functional effects of these compounds were assessed by measuring the change in the intrinsic Trp fluorescence of G_tα_. First, opsin membranes at 50 nM concentration in a buffer consisting of 20 mM BTP, 120 mM NaCl, and 1 mM MgCl_2_, pH 7.0 were incubated with a compound added to a final concentration of 10 μM for 30 min at RT, followed by pigment regeneration with 9-*cis*-retinal at 5 μM for 10 min at RT. Next, Gt was added to 500 nM concentration and the sample was illuminated for 1 min with a Fiber-Light illuminator (Dolan Jenner Industries Inc., Boxborough, MA) through a 480-520 nm bandpass wavelength filter (Chroma Technology Corporation, Bellows Falls, VT). This step was followed by the addition of 10 μM GTPγS. Measurements were performed for 1200 s with an FL 6500 Fluorescence Spectrometer. Excitation and emission wavelengths were set at 300 nm and 345 nm, respectively (18,43). Gt activation rates were determined for the first 500 s. Each condition was performed in triplicate and the experiment was repeated.

### Cell Culture

The NIH-3T3, HEK-293, 661W, and ARPE cells were cultured in DMEM with 10% fetal bovine serum (FBS) (Hyclone, Logan, UT), and 1 unit/ml penicillin with 1 μg/ml streptomycin (Life Technologies) at 37 °C under 5% CO_2_ according to the instructions from the ATCC Animal Cell Culture Guide. For NIH-3T3 DMEM lacking sodium pyruvate was used. Murine photoreceptor-derived 661W cells were provided by Dr. Muayyad Al-Ubaidi, University of Houston.

### Cytotoxicity Assay

Cells were plated in the 96-well plates at a density of 30,000 cells/ well. The next day, these cells were treated with different concentrations of the new compounds for 24 h. The cell viability was evaluated by using the 3-(4,5-Dimethyl-2-thiazolyl)-2,5-diphenyl-2H-tetrazolium bromide (MTT) cell proliferation assay (Sigma). Non-treated cells were used as control. Cytotoxicity was determined by calculating the percentage of dead cells in each experimental condition. All experimental conditions were performed in triplicate and the experiments were repeated three times.

### cAMP Detection Assay

The NIH-3T3 cells stably expressing either WT or P23H rod opsin were plated in two 96-well plates at a density of 50,000 cells/ well in 85 μl of DMEM medium containing 10% FBS and antibiotics. The cells were treated with the compounds at different concentrations for 16 h. Next, 9-*cis*-retinal at 5 μM was added for 2 h and kept in the dark to regenerate isoRho in these cells. While one plate was kept in the dark, the second plate was exposed to bright light for 15 min from a 10 cm distance. Levels of accumulated cAMP were detected with the cAMP-Glo™ kit (Promega) following the manufacturer’s protocol. The luminescence signal was recorded with a FlexStation 3 plate reader (Molecular Devices). A standard curve was developed using cAMP provided by the kit and the percentage of cAMP was calculated for each condition. The values were expressed as a percentage, assuming the cAMP level detected in the non-treated cells as 100%. Each condition was performed in triplicate and the experiment was repeated.

### Detection of Membrane Localization of Rod Opsin in NIH-3T3 cells

The NIH-3T3 cells stably expressing P23H rod opsin were plated in a 96-well plate at a density of 30,000 cells/ well and cultured for 8 h before 10 μM CR4, CR5 or 5 μM 9-*cis*-retinal were added to these cells. The plate was covered with aluminum foil and incubated overnight for 16 h. Next day, medium was removed and cells were washed 3 times with PBS and then fixed in 4% freshly prepared paraformaldehyde in PBS for 20 min in RT. Then, cells were incubated in 10% normal goat serum in PBS for 1 h at 37 °C. To detect opsin expressed on the cell surface, cells were incubated with the mouse B6-30 anti-Rho antibody, recognizing the N-terminal epitope, for 3 h at RT. Then, cells were washed 3 times with PBS for 5 min at RT. Next, cells were incubated with anti-mouse secondary antibody conjugated with Alexa Fluor 594 (Thermo Fisher Scientific) diluted 1:500 in PBS for 1 h at RT. After that, cells were washed 3 times in PBS for 5 min at RT. The cell nuclei were stained with DAPI, following the manufacturer’s protocol. The plate was sealed with a transparent film and cells were imaged with the Operetta High Content Imager (PerkinElmer Life Sciences) using a 20x objective. Nine fields were taken of each well for cell images with four channels, including bright field, GFP, Alexa Fluor 594 and DAPI. Images were analyzed with the Columbus storage and analysis system (PerkinElmer Life Sciences). DAPI fluorescence was used to visualize nuclei and count cells. These cells also express GFP. Thus, bright filed and GFP fluorescence were used to define intact cells. The plasma membrane was outlined within ±5% of the cell border. Each condition was performed in triplicate and the experiment was repeated.

### Deep Mutational Scanning

Deep mutational scanning was used to quantitatively compare the effects of CR4 and CR5 compounds on the plasma membrane expression of 123 known retinopathy variants as previously described (31). Briefly, site-directed mutagenesis was used to generate 123 individual retinopathy variants in the context of a barcoded vector containing a cDNA encoding rod opsin with an N-terminal hemagglutinin (HA) affinity tag, which was flanked by a 5’ attB recombination site and a 3’ IRES-dasherGFP expression reporter. Sanger sequencing was used to validate each mutant and match it to its corresponding unique 10 nucleotide identifier barcode within the plasmid backbone. We then used a pool of these plasmids to generate a mix of recombinant HEK-293T cells that each express a single variant from a common genomic recombination site as was previously described (30,44). We then used fluorescence-activated cell sorting (FACS) to isolate recombinant cells based on their gain in GFP expression. These recombinant cells expressing individual rod opsin variants were then surface immunostained with a Dylight 550-labeled anti-HA antibody (ThermoFisher) and then fractionated according to the immunostaining intensity of expressed variants using FACS. Cellular isolates were then expanded prior to the extraction of the recombined genomic DNA from each fraction. Illumina sequencing of the recombined barcodes was then used to infer the immunostaining intensity of each corresponding variant in the presence and absence of each compound or 9-*cis*-retinal as a control. The analysis was repeated in triplicate for each condition.

### Animals Care and Treatment Conditions

*Abca4^-/-^Rdh8^-/-^* mice were used to examine the effects of the CR5 compound against light-induced retinal degeneration. These mice were genotyped to confirm the absence of the *Rd8* mutation, but they have the Leu variation at amino acid 450 of retinal pigment epithelium 65 kDa protein (RPE65) (35,45). Heterozygous *Rho^P23H/+^ knock-in* mice were used to evaluate the effectiveness of the CR5 compound in RP. *Rho^P23H/P23H^* (Research Resource Identifier, **RRID**:IMSR_JAX:017628) were crossed with C57BL/6J mice **RRID**:IMSR_JAX:000664), Jackson Laboratory, Bar Harbor, ME) to obtain heterozygous *Rho^P23H/+^* mice. The compound dissolved in 50% DMSO in phosphate-buffered saline (PBS) was delivered to mice via intraperitoneal (i.p.) injection. Both male and female mice were used in all experiments. All mice were housed in the Animal Resource Center at the School of Medicine, Case Western Reserve University (CWRU), and maintained in a 12-hour light/dark cycle. All animal procedures and experimental protocols were approved by the Institutional Animal Care and Use Committee at CWRU and conformed to recommendations of both the American Veterinary Medical Association Panel on Euthanasia and the Association for Research in Vision and Ophthalmology as well as the National Eye Institute Animal Care and Use Committee (NEI-ASP 682). Efforts were taken to minimize animal suffering.

### Light-Induced Retinal Degeneration

Four to six weeks old *Abca4^-/-^Rdh8^-/-^* mice were dark-adapted 24 h before the treatment. The CR5 compound at a concentration of 100 mg/kg body weight (b.w.) or DMSO vehicle were administered to mice 30 min before illumination with 10,000 lux light (150-W bulb, Hampton Bay; Home Depot, Atlanta, GA) for 30 min (20,35). Before the exposure to light mice, pupils were dilated with 1% tropicamide. After light exposure, mice were kept in the dark for 7 days and then analyzed. Retinal structure was visualized *in vivo* by spectral domain-optical coherence tomography (SD-OCT) (n = 6 mice per group) and scanning laser ophthalmoscopy (SLO) (n = 6 mice per group). Retinal function was examined with electroretinography (ERG) (n = 5 mice per group). Before each procedure, mice were anesthetized with a cocktail containing ketamine (20 mg/ml) and xylazine (1.75 mg/ml) at a dose of 4 μl/g b.w. Mice were euthanized by cervical dislocation under deep anesthesia before collection of eyes. Eyes (n = 6 mice per group) were fixed for preparation of paraffin sections stained with hematoxylin and eosin (H&E).

### Treatment of *Rho^P23H/+^* mice

*Rho^P23H/+^* mice (34) were used to examine the effects of the CR5 compound on the progression of retinal degeneration in RP at 21–33 days of age as described in (19). CR5 at 10 mg/kg b.w. or vehicle were administered i.p. to mice every other day at the same time of the day. Six injections total were performed. Retinal morphology was visualized with SD-OCT (n = 6 mice per group) and retinal function was examined with ERG (n = 5 mice per group). Before each procedure, mice were anesthetized with a cocktail containing ketamine (20 mg/ml) and xylazine (1.75 mg/ml) at a dose of 4 μl/g b.w. Mice were euthanized by cervical dislocation under deep anesthesia before eyes collection (n= 6 per group) for histological evaluation and immunohistochemistry.

### SD-OCT

The ultrahigh-resolution Spectral-Domain Optical Coherence Tomography (SD-OCT) (Bioptigen, Morrisville, NC) *in vivo* imaging of the retina was used to examine the protective effect of the CR5 compound against retinal degeneration induced by a bright light in *Abca4^-/-^Rdh8^-/-^* (46) or spontaneous photoreceptor degeneration caused by an inherited mutation in *Rho^P23H/+^* mice (34). The a-scan/b-scan ratio was set at 1200 lines. The OCT images were obtained by scanning at 0 and 90 degrees in the b-mode. Five image frames were captured and averaged. Changes in the retinas of treated and control mice were determined by measuring the thickness of the outer nuclear layer (ONL) at 0.5 mm from the optic nerve head (ONH). Six mice were used in each experimental group.

### SLO

The Scanning Laser Ophthalmoscopy (SLO) (Heidelberg Engineering, Franklin, MA) was used for the *in vivo* whole-fundus imaging of mouse retinas. Immediately after the performed SD-OCT imaging, *Abca4^-/-^Rdh8^-/-^* mice were subjected to the SLO imaging using the auto-fluorescence mode. The number of autofluorescent spots (AF) detected in different experimental groups was quantified and compared to determine the statistical significance. Six mice were used in each experimental group.

### Retinal Histology

Mouse eyes (n = 6 per group) were collected from euthanized mice and fixed in 0.5% glutaraldehyde in 2% paraformaldehyde (PFA) in PBS for 24 h at RT on a rocking platform followed by their incubation in 1% PFA for 48 h at RT. These eyes were embedded in paraffin and sectioned. Sections (5 μm thick) were stained with H&E. The retina imaging was performed with a ZEISS Axio Scan.Z1 slide scanner (Carl Zeiss Microscopy GmBH, Jena, Germany) and analyzed using Zeiss-Zen 3.2 software (blue edition).

### Immunohistochemistry

To detect the rod and cone photoreceptors in the retina, eyes were collected from CR5-treated or vehicle-treated euthanized *Rho^P23H/+^* mice at P33 fixed in 4% PFA for 24 h followed by their incubation in 1% PFA for 48 h at RT. The eight-μm thick cryo-sections prepared from fixed eyes were blocked with 10% normal goat serum (NGS) and 0.3% Triton X-100 in PBS for 1 h at RT and then stained overnight at 4°C with a monoclonal mouse 1D4 anti-Rho primary antibody to visualize rod photoreceptors and biotinylated peanut agglutinin (PNA) (1:500 dilution) to visualize cone photoreceptors. The next day sections were washed with PBS, followed by the incubation with Alexa Fluor 555-conjugated goat anti-mouse secondary antibody (1:400 dilution) to detect rods and Fluor 488-conjugated streptavidin (1:500 dilution) to detect cones for 2 h at RT. Cell nuclei were detected by staining with DAPI. Slides were coverslipped with Fluoromount-G (SouthernBiotech).

### ERG

The electroretinography (ERG) was used to assess the effect of the CR5 compound on retinal function in *Abca4^-/-^Rdh8^-/-^* mice injured with bright light and *Rho^P23H/+^* mice, modeling human RP. Scotopic and photopic ERGs were recorded for both eyes of each mouse using a Celeris rodent ERG system and Espion Dyagnosys software Version 6 (Dyagnosys, LLC, Lowell, MA). The ERG data were processed for each experimental group and presented as mean and standard deviation (S.D.) for both a-wave and b-wave amplitudes. Each experimental group contained five mice.

### Statistical Analyses

The compound cytotoxicity, thermal shift assay, and quantification of cAMP levels were performed in triplicates and repeated two times. Each compound was tested at different concentrations. Each assay had included positive and negative controls. The opsin-ligand binding, pigment regeneration, thermal stability, and Gt activation assays were performed three times. The effect of each compound was shown in a dose-dependent or time-dependent manner. The parameters derived from these measurements were shown as an average and S.D. For multiple comparisons, one or two-way ANOVA with Turkey’s post hoc tests were used. All statistical calculations were performed using the Prism GraphPad 7.02 software. Type1 error tolerance for the experiments was established at 5%. Values of *P* < 0.05 were considered statistically significant. Collection of data and its statistical analysis were carried out by distinct personnel.

## RESULTS

### Discovery of novel natural product-like compounds targeting the opsin’s orthosteric site

To identify novel pharmacological chaperones of rod opsin we performed a virtual screening of the natural product-derived synthetic molecules from the OTAVA collection (composed of 1547 compounds) and used the crystal structure of bovine rod opsin (PDB ID: 3CAP) as a target. The compounds were docked to the opsin’s chromophore-binding pocket using VINA/Vega 3.1.0.21 software to calculate the binding free energies (36). Quercetin was used as a positive control and the value of the binding free energy obtained for quercetin-rod opsin complex (−9.3 kcal/mol) was used as a cut-off to select the hit compounds. Of the 100 top compounds identified in the initial screening twenty five compounds with binding free energy lower than that obtained for quercetin were re-scored. These compounds were suitable for an evaluation to assess the position into the orthosteric site and coverage of different chemical scaffolds, avoiding overflowing the library with similar chemical compounds. This analysis resulted in the identification of five hit compounds (CR1-CR5) with different binding patterns and chemical scaffolds shown in the Tanimoto plot (**Fig. 1A and B**). All these compounds contained a common chromenone motif. These compounds bound to rod opsin orthosteric site with the following binding free energies, CR1: −11.3 kcal/mol, CR2: −10.5 kcal/mol, CR3: −10.0 kcal/mol, CR4: −10.3 kcal/mol, CR5: −10.3 kcal/mol). The positions of these compounds within the chromophore-biding pocket are shown in **Fig. 1C**. All five compounds accommodated within the retinal-binding pocket in the proximity to the active Lys296 residue and other residues involved in the stabilization of the native chromophore 11-*cis*-retinal, including Ala117 and Ala292. All these compounds formed hydrogen bond interactions with Glu181 and Tyr191, two residues found to coordinate quercetin within the orthosteric binding site. The details of the compound-opsin interactions obtained by docking analyses are shown in **Fig. S1**.

**Figure 1.**
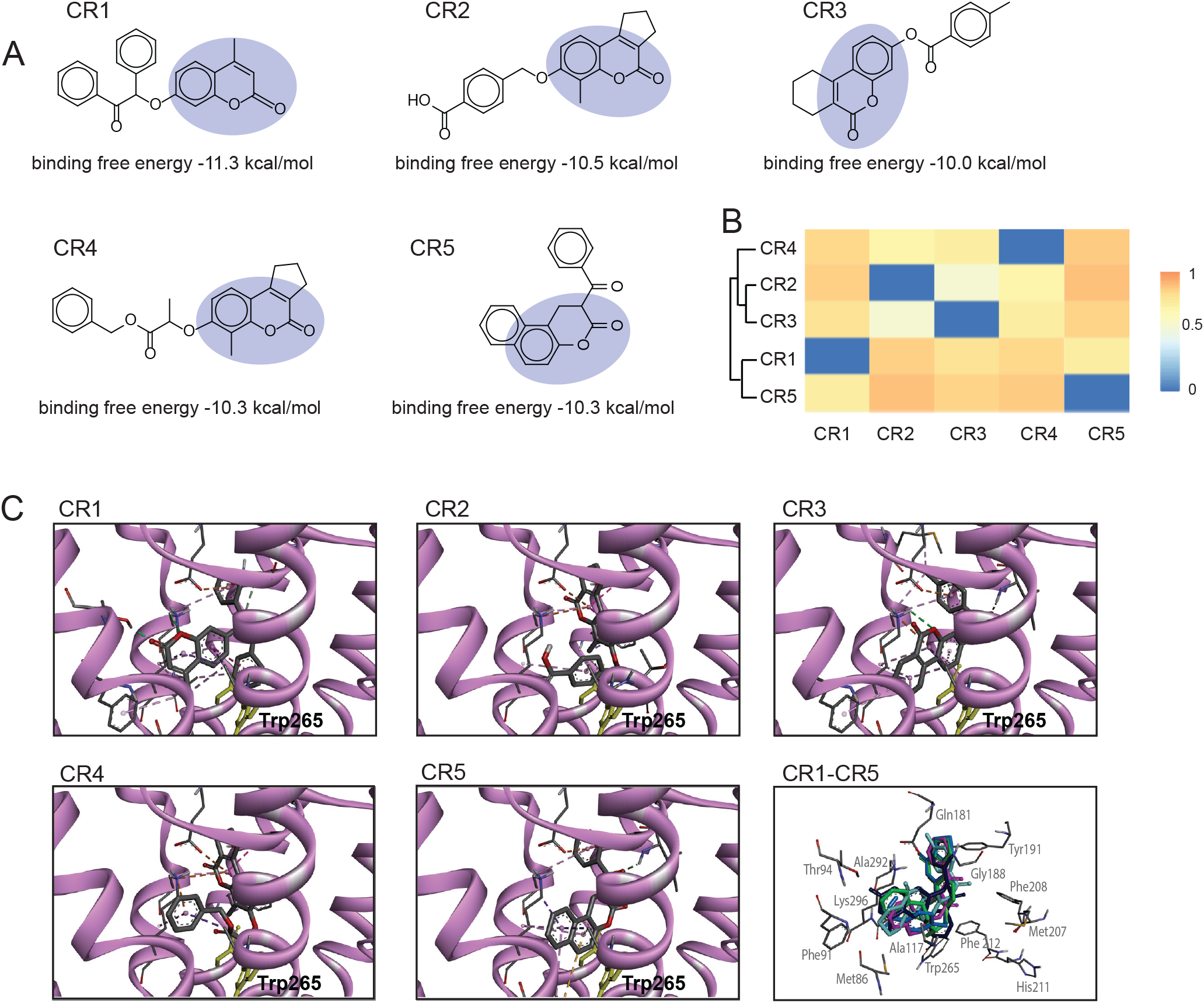
Rod opsin modulators with a natural product scaffold identified from the virtual screening. Five compounds that emerged from *in silico* analysis were purchased from OTAVAchemicals Ltd. for further evaluation. **A**, Chemical structures of these compounds and binding free energies are shown. The chromenone motif is highlighted. **B**, Hierarchical clustering of identified compounds. Hierarchical clustering of the hit compounds was built based on pairwise compound similarities using atom pair description and Tanimoto coefficient. The structural differences are depicted as a heat map to show the distance matrix between the five compounds. **C**, Molecular docking of the new chromenone ligands to the orthosteric site of rod opsin. The best binding poses of five compounds identified *in silico* are shown within the orthosteric binding pocket of bovine rod opsin (PDB ID: 3CAP). Rod opsin structure is shown in violet. The carbon atoms of the compounds are shown in grey. The overlay of all five compounds within the opsin’s orthosteric site is also shown (CR1, dark blue; CR2, blue; CR3, light blue; CR4, green; CR5, purple). Structures obtained from molecular docking simulations were visualized with the Biovia Discovery Studio Visualizer 17.2.0 visualizer.

### Evaluation of drug-likeness properties of the chromenone lead compounds

To determine whether the identified chromenone compounds exhibit drug-like properties, we analyzed their absorption, distribution, metabolism, and elimination (ADME) profiles by using the SWISSADME server. The main ADME parameters, including lipophilicity, hydrophilicity, solubility, absorption in the gastrointestinal (GI) tract, permeability to the blood-brain barrier (BBB) are presented in **Table 1**. None of the selected compounds violated the drug-likeness associated with Lipinski’s rule of five. Thus, all five compounds could be potential therapeutic lead compounds based on their moderate solubility, high predicted absorption in the GI tract, and most importantly predicted ability to cross the BBB, and thus also likely the blood-retina barrier (BRB).

**Table 1.** Pharmacokinetic properties of the identified *in silico* compounds

| Molecule | MW | Lipophilicity<br>(Log P) | Hydrophilicity<br>(LogS) | Water<br>Solubility | GI<br>Absorption | BBB<br>Permeant | Lipinski<br>Violations | PAINS<br>Alerts | Synthetic<br>Accessibility |
| --- | --- | --- | --- | --- | --- | --- | --- | --- | --- |
| CR1 | 370.4 | 4.47 | -5.75 | Moderately<br>soluble | High | Yes | 0 | 0 | 3.76 |
| CR2 | 350.36 | 3.7 | -4.51 | Moderately<br>soluble | High | Yes | 0 | 0 | 3.49 |
| CR3 | 334.37 | 4.34 | -5.1 | Moderately<br>soluble | High | Yes | 0 | 0 | 3.47 |
| CR4 | 378.42 | 4.25 | -5.02 | Moderately<br>soluble | High | Yes | 0 | 0 | 4.23 |
| CR5 | 302.32 | 3.74 | -4.93 | Moderately<br>soluble | High | Yes | 0 | 0 | 2.98 |

Toxicity is another important parameter to be evaluated for the new compound. Thus, the potential cytotoxicity of the identified chromenone derivatives was examined in the NIH-3T3 and HEK-293 cells used for the *in vitro* biochemical analyses in this current study. Also, taking into consideration that these new chromenone compounds have not been evaluated in the retina-like system we tested toxicity of these compounds in two cell lines related to vision, namely photoreceptor-derived 661W cells, and ARPE19 cells, which were derived from retinyl pigment epithelial (RPE) cells (**Fig. S2**). These cells were treated overnight with CR1-CR5 compounds at a concentration range of 0.001-100 μM. No toxicity was found for up to 10 μM compound concentration. However, treatment with 100 μM compound resulted in decreased cell viability by 20-40%. Thus, overall these compounds did not exhibit relevant toxicity at concentrations below 100 μM and could be evaluated for their potential as pharmacological chaperones of RP-linked rod opsin mutants.

### Evaluation of the binding of chromenone derivatives to rod opsin

To confirm the binding of the new identified *in silico* compounds into the opsin’s orthosteric binding pocket we used tryptophan fluorescence quenching assay. In the ligand-free opsin conformation the fluorescence intensity of the Trp265 residue located within the chromophore-binding pocket reaches its maximum. However, this fluorescence is quenched after binding of the native ligand, 11-*cis*-retinal, due to the electrostatic interactions formed between the β-ionone ring of the retinal and the residue side chain. Our molecular docking analysis indicated that the binding of the identified five compounds is also likely to generate Trp265 quenching. To test this interaction, opsin within ROS discs membranes was titrated with increasing concentrations of the compounds and the intrinsic Trp fluorescence at 330 nm was monitored. Each of these compounds resulted in fluorescence quenching with an EC_50_ at low nanomolar concentrations (CR1 EC_50_ = 275 ± 26 nM, CR2 EC_50_ = 375 ± 21 nM, CR3 EC_50_ = 207 ± 46 nM, CR4 EC_50_ = 171 ± 23 nM, CR5 EC_50_ = 193 ± 40 nM). The EC50 values obtained for CR1-CR5 compounds were similar to those reported for 9-*cis*-retinal analog (EC_50_ = 124 nM to 1.2 μM) (16,47). These results indicate that the new chromenone compounds can accommodate within the opsin’s orthosteric site (**Fig. 2A**).

**Figure 2.**
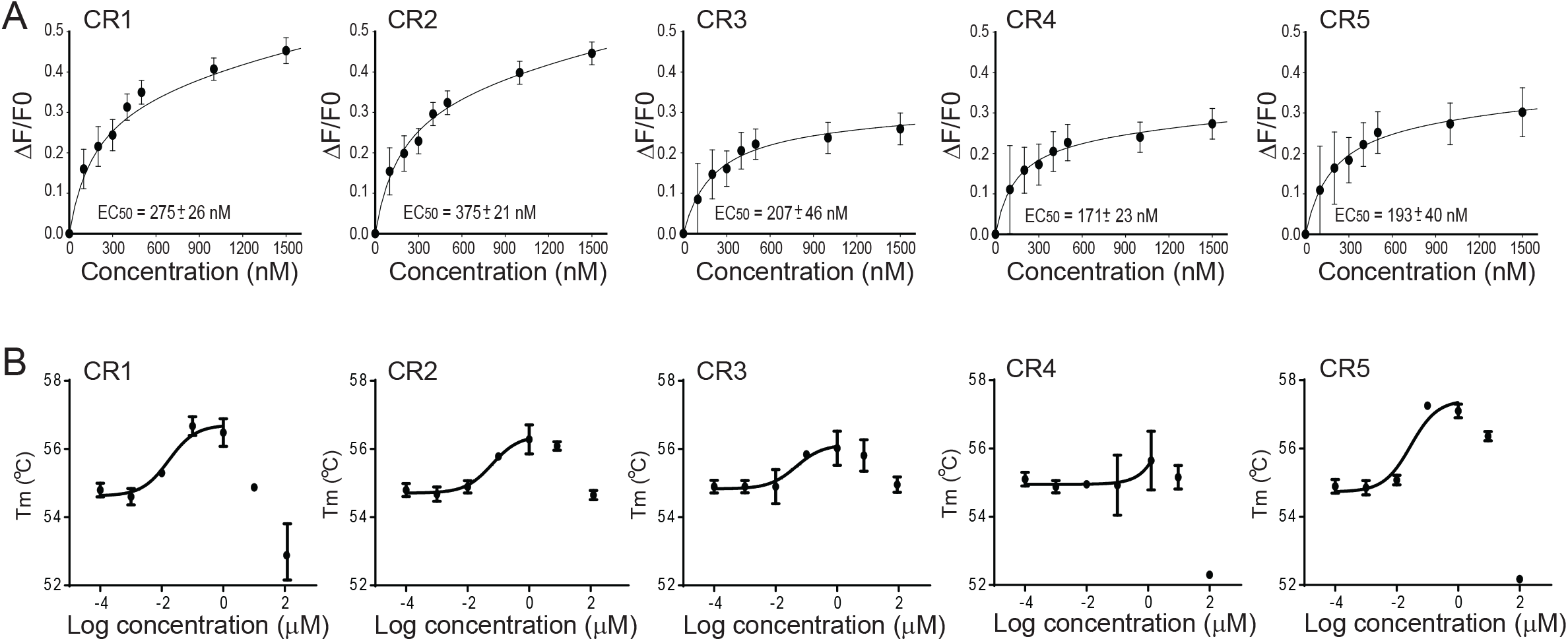
Binding of the new chromenone ligands to the opsin’s orthosteric site. **A**, The binding of compounds CR1-CR5 was determined by measuring the quenching of the intrinsic Trp fluorescence by these compounds added to bovine rod opsin membranes at different concentrations (100-1500 nm). Changes in the fluorescence intensity at 330 nm were plotted as a function of the compound concentration. The binding curves were fitted using PRISM GraphPad software. EC50s of each compound were calculated and averaged from triplicates. These values ± standards deviation (S.D.) are shown in the figure. **B**, Effect of the new compounds on the temperature of melting (Tm) of opsin. Tm was determined for the bovine rod opsin membranes incubated with the CR1-CR5 compounds at different concentrations (0.0001-100 μM) by using a fluorescent probe BiFC. The change in fluorescence intensity was measured in response to increasing temperature, and the Tm was plotted against the corresponding ligand concentration. Error bars represent S.D. Each measurement was performed in triplicate and the experiment was repeated three times.

The conformational change that occurs upon the accommodation of a ligand within the opsin’s orthosteric site also can be detected by measuring its temperature of melting (Tm) (16–18). Thus, we used this property to validate further binding of the newly identified compounds to opsin. The ROS membranes containing unliganded opsin were incubated with the compounds at different concentrations and then heated up to determine the temperature at which opsin denatures. Lower concentrations of these compounds (0.1-10 μM) consistently increased the Tm 1-3 °C above the Tm of unliganded opsin (Tm = 54.8±02 °C) in a dose-dependent manner (**Fig. 2B and Table 2**). Tm values in the presence of 1 μM concentration of each compound were as follows: CR1 Tm = 56.5 ± 0.4 °C, CR2 Tm = 56.3 ± 0.4 °C, CR3 Tm = 56.1 ± 0.5 °C, CR4 Tm = 55.6 ± 0.9 °C, CR5 Tm = 56.8 ± 0.6 °C. In each case, incubation with the chromenone compounds caused a modest increase in opsin’s Tm. Of note, elevated concentrations of all these molecules (≥100 μM) caused a decrease in Tm, which may reflect the non-specific effects of the drugs on the structure of the membrane and/ or protein. Nevertheless, the consistent increase in Tm caused by the lower concentrations of the chromenone compounds confirm that the binding of these ligands is coupled to rod opsin folding. Of the five compounds tested, the most pronounced effect on the opsin’s Tm was produced by CR5 (at 0.1 μM concentration Tm = 57.3 ± 0.6 °C). The observed changes in the rod opsin’s Tm upon incubation with the chromenone compounds confirm that these molecules bind to unliganded opsin and modulate its conformation.

**Table 2.**
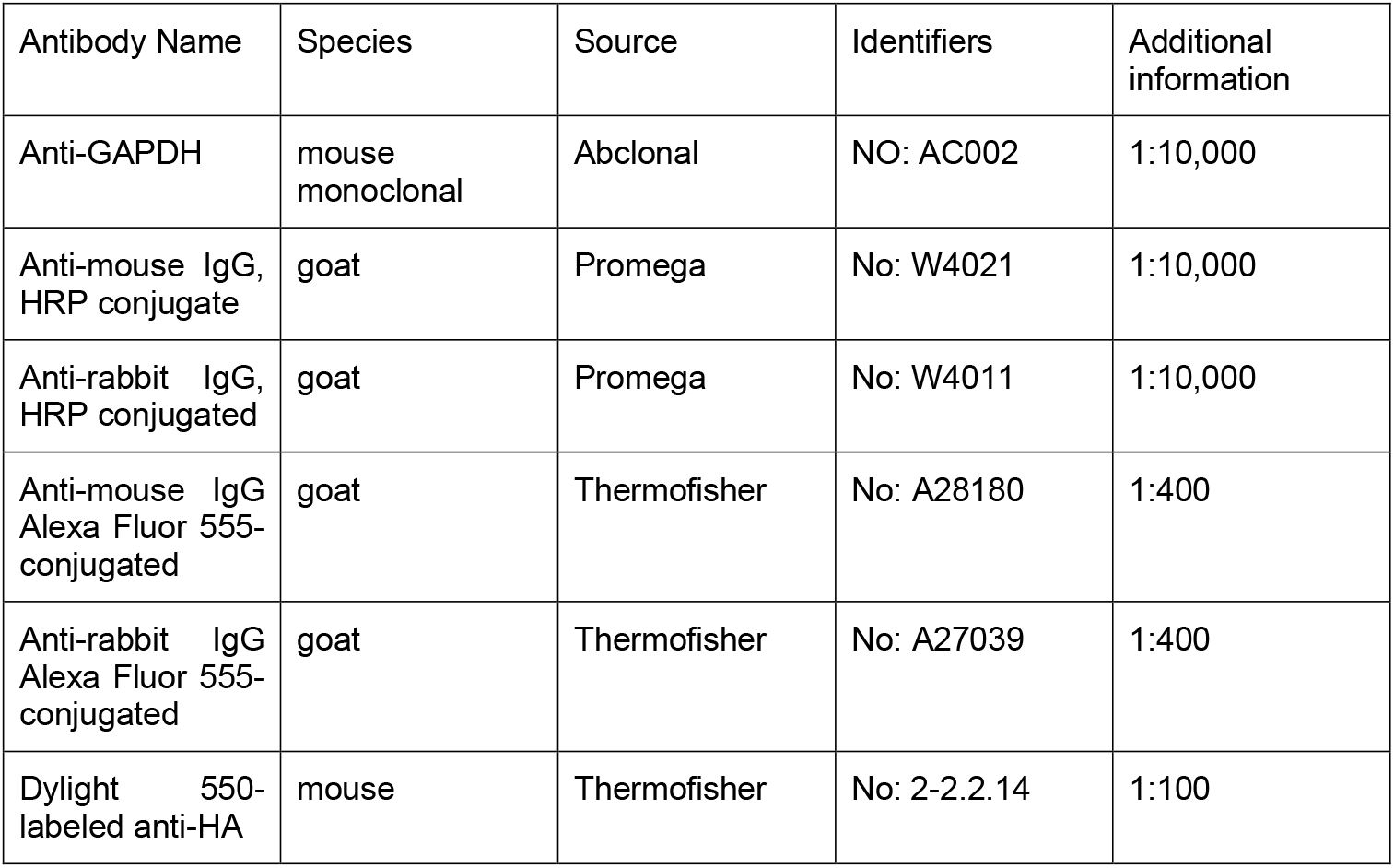
The list of used commercial antibodies.

**Table 2.** The temperature of melting of rod opsin in the presence of the new compounds

| Concentration<br>(μM) | T<sub>m</sub> | | | | |
| --- | --- | --- | --- | --- | --- |
|  | CR1 | CR2 | CR3 | CR4 | CR5 |
| **0.0001** | 54.8 ± 0.2 | 54.8 ± 0.2 | 54.9 ± 0.2 | 55.1 ± 0.2 | 54.9 ± 0.2 |
| **0.001** | 54.6 ± 0.2 | 54.7 ± 0.2 | 54.9 ± 0.2 | 54.9 ± 0.2 | 54.9 ± 0.2 |
| **0.01** | 55.3 ± 0.1 | 54.9 ± 0.2 | 54.9 ± 0.2 | 55.0 ± 0.08 | 55.1 ± 0.1 |
| **0.1** | 56.7 ± 0.3 | 55.8 ± 0.02 | 55.8 ± 0.1 | 54.9 ± 0.9 | 57.3 ± 0.1 |
| **1** | 56.5 ± 0.4 | 56.3 ± 0.4 | 56.1 ± 0.5 | 55.6 ± 0.9 | 56.8 ± 0.6 |
| **10** | 54.9 ± 0.04 | 56.2 ± 0.2 | 55.8 ± 0.7 | 55.2 ± 0.6 | 56.0 ± 0.2 |
| **100** | 53.3 ± 1.4 | 54.3 ± 0.2 | 54.9 ± 0.4 | 51.8 ± 0.02 | 51.8 ± 0.02 |

### Effect of the compounds on the pigment regeneration

To examine if the identified compounds affect the retinal binding to opsin, we monitored the changes in the UV-visible absorbance spectra upon addition of 9-*cis*-retinal to ROS membranes preincubated with the compound and compared to the absorbance spectrum obtained for opsin incubated with 9-*cis*-retinal alone. The binding of retinal analogs could be described as a dynamic process occurring in two stages, the early stage related to the initial interaction between the ligand and the receptor and a late-stage associated with the formation of the covalent Schiff base linkage, which results in the appearance of the maximum absorption peak at 485 nm. Although the retinal channeling is not fully understood, the structural and biochemical evidence indicates that the retinal entry site is located between transmembrane helices TM5 and TM6, while the exit site is situated on the opposite side of the channel between TM1 and TM7 (48–50). The interaction of retinal and opsin in the early stage of the entering process induces a conformational rearrangement in the binding pocket, which dynamically adapts to facilitate the entry and stabilization of the incoming ligand. Thus, for non-retinoid ligands bound to opsin through non-covalent interactions, these structural changes induced by the retinoid could become a major destabilization factor. This destabilization likely disrupts the interactions between the non-retinoid compound and the residues within the binding pocket, resulting in the compound release through the exit site. Three (CR1-CR3) of five compounds did not affect the formation of the isoRho pigment as evidenced by no change in the absorption maximum at 485 nm, suggesting their weak binding within the orthosteric site that are displaced by incoming retinal. On the contrary, the CR4 and CR5 compounds affected pigment formation. Preincubation of opsin with CR4 resulted in a slight increase in the λ_max_ of isoRho. Conversely, CR5 decreased the total pigment formation by ~20.5±7.4%. These results suggest that CR4 and CR5 bind to opsin more favorable than CR1-CR3. Indeed, molecular dynamics simulations of CR4 and CR5 bound states suggest the association of these molecules locks opsin into native-like conformation (**Fig. S3**). However, different effects of CR4 and CR5 on the isoRho formation suggests that these two compounds likely accommodate within the orthosteric site differently (**Fig. 3A**).

**Figure 3.**
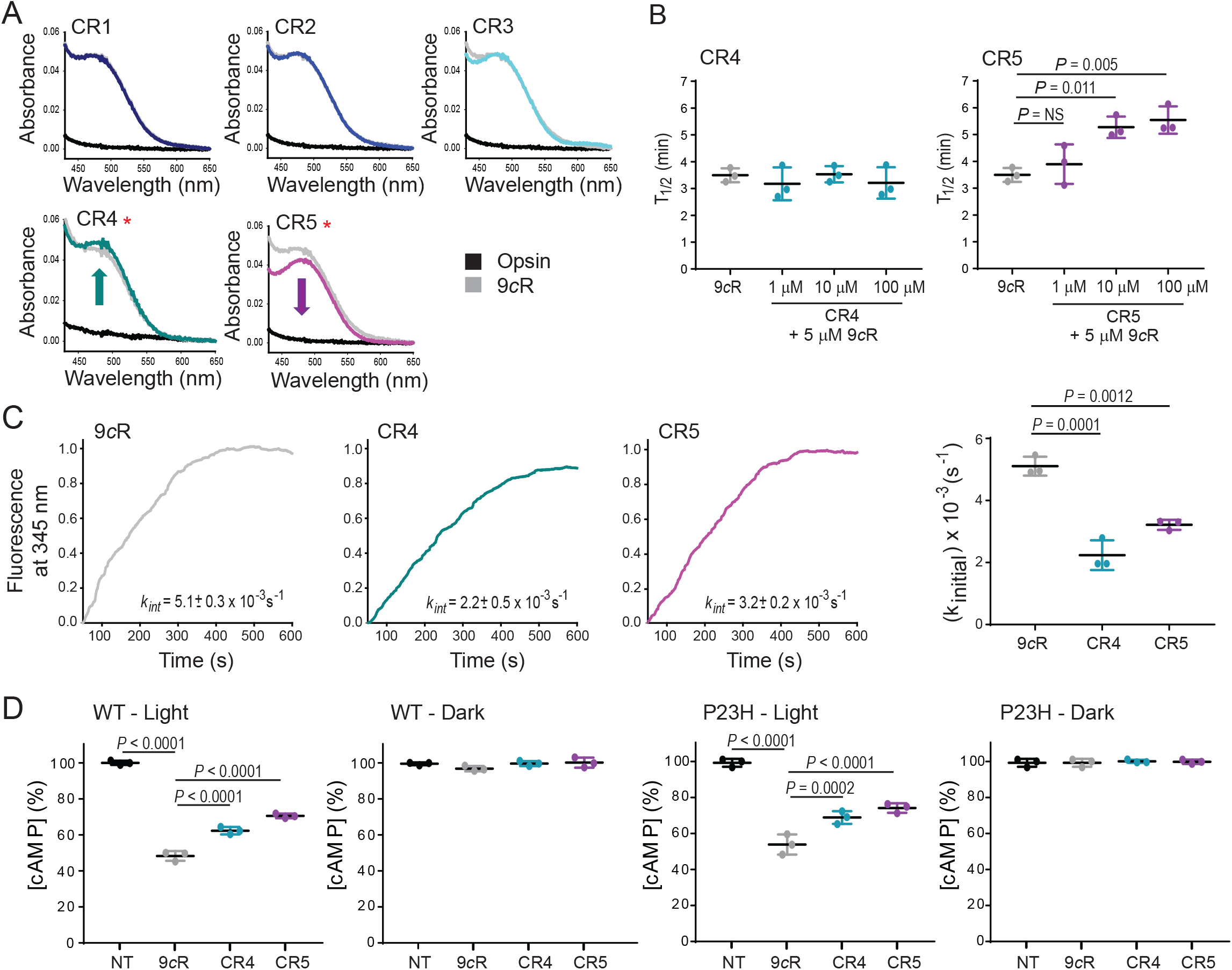
Effects of the new chromenone derivatives on pigment regeneration and function. **A**, UV-visible spectra of isoRho regenerated with 5 μM 9-*cis*-retinal (9*c*R) after treatment of opsin membranes with the CR1-CR5 compounds at 10 μM concentration. **B**, Half-time (T1/2) of isoRho regenerated with 5 μM 9*c*R after treatment of bovine rod opsin membranes with the CR4 and CR5 compounds at 1, 10 and 100 μM concentrations. Error bars represent the standard deviation (S.D.). Each measurement was performed in triplicate and the experiment was repeated. Statistical analysis was performed with the one-way ANOVA and Turkey post hoc tests. The statistically different changes (*P* < 0.05) are indicated in the figure. **C**, Gt activation by the CR4 or CR5 compound-bound isoRho within membranes. Bovine rod opsin membranes were incubated with 10 μM compound prior to regeneration with 5 μM 9-*cis*-retinal (9*c*R). The activity of illuminated isoRho was recorded by monitoring changes in the Trp fluorescence at 345 nm related to the dissociation of the isoRho-Gt complex upon the addition of 10 μM GTPγS. These changes were plotted as a function of time. Excitation and emission wavelengths were 300 and 345 nm, respectively. Initial rates and error bars (S.D.) for each condition were plotted and shown in the right panel. Each measurement was performed in triplicate and the experiment was repeated. **D**, Effect of CR4 and CR5 on WT and P23H isoRho function in cultured cells. Rho can couple to Gi signaling within cells and modulate the concentration of cellular levels of cAMP upon its activation. Levels of cAMP in the NIH-3T3 cells stably expressing WT or P23H rod opsin regenerated with 5 μM 9*c*R were monitored either in the dark or upon light illumination. Cells were treated with CR4 or CR5 compound overnight before isoRho regeneration. Cells treated with 9*c*R only and nontreated (NT) cells were used as controls. Each condition was performed in triplicate and the experiment was repeated. Statistical analysis was performed with the one-way ANOVA and Turkey’s post hoc tests. The statistically different changes (*P* < 0.05) are indicated in the figure.

Based on the cumulative results obtained from these biochemical analyses, including Trp fluorescence quenching, effect on Tm, and pigment regeneration, we chose to characterize the effects of CR4 and CR5 in further detail. First, we measured the effect of each compound on the rate of pigment regeneration by monitoring the time-dependent change in the absorbance of the pigment in the presence of these compounds at 3 different concentrations (1, 10, and 100 μM) incubated with opsin prior to the addition of 5 μM 9-*cis*-retinal (**Fig. 3B, Table 3**). Only the CR5 compound slowed the kinetics of pigment regeneration as evidenced by the increase in the apparent half-time (t_1/2_) (t_1/2_ = 5.3 ± 0.4 min at 10 μM and t_1/2_ = 5.5 ± 0.5 min at 100 μM) as compared to t_1/2_ = 3.9 ± 0.6 min obtained for 9-*cis*-retinal. The pigment regeneration kinetics in the presence of CR4 was slightly enhanced (t_1/2_ = 3.5 ± 0.3 min at 10 μM and t_1/2_ = 3.2 ± 0.6 min at 100 μM) as compared to 9-*cis*-retinal alone, though the observed changes were not significant. Nevertheless, these results suggest that CR4 and CR5 modulate opsin by altering the retinal chromophore entry in a different manner.

**Table 3.** IsoRho pigment regeneration rates and its stability at 55 °C regenerated in the presence and in the absence of CR4 and CR5.

| Concentration<br>($\mu\text{M}$) | $T_{1/2}$ (min) | | |
| --- | --- | --- | --- |
|  | 9cR | CR4 | CR5 |
| **1** | - | $3.2 \pm 0.6$ | $3.9 \pm 0.7$ |
| **5** | $3.5 \pm 0.2$ | - | - |
| **10** | - | $3.5 \pm 0.3$ | $5.3 \pm 0.4$ |
| **100** | - | $3.2 \pm 0.6$ | $5.5 \pm 0.5$ |

### Functional effects of top compounds on rod pigment

To assess the functional effects of CR4 and CR5 on the visual pigment, we measured the activation rates of its cognate G protein, G_t_. Bovine rod opsin membranes were pre-incubated with each compound for 15 min prior to the regeneration of the pigment with 9-*cis*-retinal for 10 min. We then added purified G_t_ and illuminated the mixture to activate the receptor and initiate the isoRho-G_t_ complex formation followed by its dissociation upon addition of GTPγS, which was detected as a change in a fluorescence arising from the dissociation of G_tα_. Both compounds decreased the initial rates of Gt activation, CR4 (*k_int_* = 2.2 ± 0.5 × 10^-3^ s^-1^) by 2.3 fold and CR5 (*k_int_* = 3.2 ± 0.2 × 10^-3^ s^-1^) by 1.6 fold as compared to isoRho (*k_int_* = 5.1 ± 0.3 × 10^-3^ s^-1^), indicating that both compounds are capable of modulating the function of the rod visual pigment *in vitro* (**Fig. 3C**).

Although Gt is specifically expressed in the retina to mediate signaling from the activated rod visual receptors, heterologously expressed rod pigment can signal through G_i_ present in mammalian cells. To determine if binding of either of these compounds modulate downstream signaling, we measured the effects of the CR4 and CR5 compounds on cAMP levels in the NIH-3T3 cells stably expressing rod opsin. Cells were treated with CR4 or CR5 at 10 μM concentration overnight followed by the regeneration with 5 μM 9-*cis*-retinal for 2 h on the next day. The effects of these compounds on cAMP accumulation were analyzed upon light stimulation (**Fig. 3D**). The levels of cAMP were reduced substantially in the WT opsin-expressing cells treated with 9-*cis*-retinal and exposed to light as compared to the non-treated control cells. However, the levels of cAMP were higher in cells treated with CR4 or CR5, followed by 9-*cis*-retinal as compared to cells treated with 9-*cis*-retinal only, indicating the modulatory effects of these compounds for the rod pigment function and confirming our *in vitro* results. As expected, treatment with CR4 and CR5 did not affect cAMP levels in the dark. Interestingly, similar results were obtained in the cells expressing the P23H rod opsin mutant (**Fig. 3D**).

Together, our results emerging from the computational analysis, biochemical binding assays, and functional examinations indicate that CR4 and CR5 compounds could modulate the Rho signaling through direct interactions with the receptor.

### Effect of the compounds on the rod opsin expression and membrane targeting of P23H mutant

To determine if the new chromenone rod opsin ligands can improve folding and rescue membrane targeting of the RP-linked mutants we used the NIH-3T3 cells stably expressing the P23H variant. As shown before, the P23H mutation results in the ER retention of this receptor, while binding of 9-*cis*-retinal analog induces its proper transport to the plasma membrane (5–7). In this study, high-content imaging was used to measure the plasma membrane expression of rod opsin in cells treated with either 10 μM CR4 or CR5 relative to untreated and 5 μM 9-*cis*-retinal-treated cells (**Fig. 4A**). Our results showed that treatment with these new compounds induced the movement of P23H rod opsin to the cell surface. The quantification of the cell surface expression indicated a considerable increase in the membrane localization of P23H rod opsin upon treatment with these compounds (**Fig. 4B**). As noted, the effect of CR5 was greater than CR4. However, these compounds had a lower effect than 9-*cis*-retinal, which is possibly related to their distinct interaction pattern and inability to form a covalent Schiff base bond with opsin. The increase in membrane expression of P23H rod opsin was not related to the increase in this receptor expression. The SDS-polyacrylamide gel and immunoblotting analyses indicated that CR4 and CR5 did not change the total expression levels of WT or the P23H rod opsin mutant, indicating that the increased level of the mutant in the plasma membrane resulted from the enhanced membrane trafficking of this receptor (**Fig. 4C**). In addition, the ratio of mature receptor (~75 kDa protein band) to immature receptor (~37 kDa protein band) increased about 1.2 ± 0.1 and 1.3 ± 0.1 fold upon treatment with CR4 and CR5, respectively, indicating their positive effect on the mutant receptor folding. These positive chaperone-like effects of CR4 and CR5 compounds encouraged the question of what other pathogenic Rho mutants could be corrected by these compounds.

**Figure 4.**
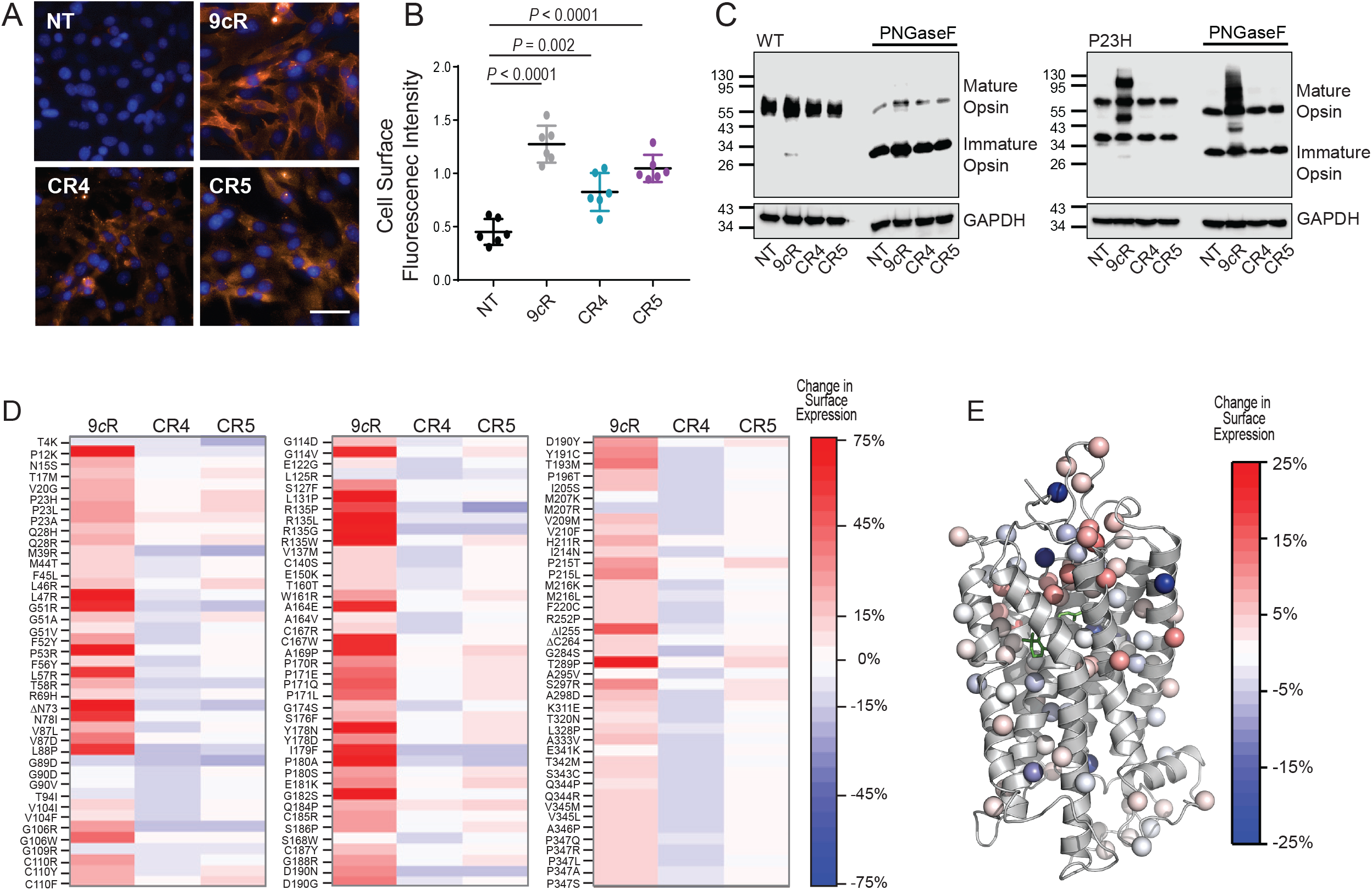
Effect of the new chromenone compounds on membrane localization of RP-related rod opsin mutants. **A**, Fluorescence images of the NIH-3T3 cells stable expressing P23H rod opsin treated either with CR4, CR5 at a final concentration of 10 μM or 5 μM 9-*cis*-retinal (9*c*R) for 16 h. The images were taken with a high-content imaging operetta microscope at 20x magnification. A 50 μm scale bare is shown for reference. The cell surface expression of opsin appears in orange. The nuclei of the cells are shown in blue. **B**, The cell surface fluorescence intensity was quantified as is described in the Material and Methods. Each condition was performed in triplicate and the experiment was repeated. Statistical analysis was performed with the one-way ANOVA and Turkey’s post hoc tests. The statistically different changes (*P* < 0.05) are indicated in the figure. **C**, Immunoblot showing the effect of CR4 and CR5 compounds on the expression level of WT and P23H rod opsin in the NIH-3T3 cells stably expressing these receptors. Total cell extracts (50 μg) were loaded and separated using the SDS-PAGE gel, followed by transfer to polyvinyl difluoride membrane (PVDF). Rod opsin was detected with the 1D4 anti-Rho antibody recognizing the C-terminal epitope of this receptor. Anti-GAPDH antibody was used as a loading control. PNGaseF-treated samples were deglycosylated prior to loading onto the gel. The experiment was repeated three times. Representative immunoblots are shown. **D-E**, Deep mutational scanning was used to measure the change in the plasma membrane expression of 123 Rho variants containing 119 adRP variants and 4 CSNB variants upon treatment with 10 μM CR4 or CR5 compounds in comparison to the treatment with 5 μM 9-*cis*-retinal (9*c*R). **D**, The heatmap depicts the surface immunostaining intensities for a collection of Rho variants bearing individual amino acid substitutions. Values represent the average from three biological replicates, and color bars indicate the scale of the observed effects under each condition. Red indicates an increase in the plasma membrane expression and blue indicates a reduction in the plasma membrane expression under each condition. **E**, The Cα of mutated side chains are rendered as spheres in the context of the three-dimensional structure of Rho (PDB ID: 1U19). Spheres are colored according to the average change in the plasma membrane expression in the presence of CR5. The 11-*cis*-retinal chromophore is shown as green sticks for reference.

### Effects of CR4 and CR5 on inherited rod opsin variants

To learn if CR4 and CR5 compounds could promote the correct transport of other disease-related rod opsin mutants, we used a recently reported deep mutational scanning assay to survey their effects on the plasma membrane expression of 123 rod opsin variants related to RP (29–31). Of note, a few CSNB variants were included in this screening. Our results revealed a significant difference in the chaperone-like properties of CR4 and CR5. While CR4 had only a minor effect on a few mutants, CR5 enhanced the surface immunostaining of 31 out of 123 mutants (**Fig. 4D and F**). Interestingly, most of the mutated residues corrected by CR5 are involved in rod opsin stabilization and retinal binding. These mutations include five within the N-terminal cap that stabilize opsin during biogenesis (T17M, P23H, P23L, P23A, and Q28R), two within the first transmembrane helix TM1 (L46R and G51A), two that disrupt a native disulfide bond (C110Y and C110F), one that interacts with the retinal (G114V), one that interacts with Gt (R135W), six within TM4 (W161R, A169P, P170R, P171E, P171Q, and P171L), nine within a cluster of residues that form a retinal plug in the ECL2 (S176F, Y178N, Y178D, P180S, E181K, Q184P, S186P, G188R, and D190Y), one near the N-terminal tip of TM7 (G284), and three near the retinal binding site in TM7 (T289P, S297R, and A298D).

These results clearly indicate that the CR5 compound could partially rescue many RP-linked Rho variants, thus suggesting that CR5 represents a promising lead compound for the development of potent novel pharmacological chaperones for misfolded RP variants.

### Therapeutic effect of CR5 in mouse models of retinal degeneration

Our *in vitro* findings encouraged us to validate the effectiveness of the CR5 compound *in vivo*, in mouse models of retina degeneration. The sequence of human and mouse rod opsin shares 95% identity and mouse models are commonly used to study pathophysiology of human retinopathies (43). Beneficial effects of rod opsin stabilizers, including polyphenolic compounds on retinal health, have been demonstrated before in *Abca4^-/-^Rdh8^-/-^* mice susceptible to bright light insult (16,20,35). Excessive concentrations of unliganded opsin that form upon illumination contributes to retinal degeneration in these mice, which could be prevented by a single administration of opsin stabilizing compounds, including 11-*cis*-6mr-retinal analog, flavonoids, or non-retinoid opsin antagonist YC-001 prior to light (16,20,35). Similarly, treatment of *Abca4^-/-^Rdh8^-/-^* mice with CR5, a novel chomenone opsin ligand, at 100 mg/kg b.w. 30 min before bright light illumination prevented damage of photoreceptors in these mice (**Fig. 5A**). As evidenced by the *in vivo* optical coherence tomography (OCT) imaging and histological evaluation, retinas of these mice closely resembled retinas of mice kept in the dark, while mice treated with the vehicle exhibited severe retina deterioration (**Fig. 5B and C**). The outer nuclear layer (ONL), where the photoreceptor cells reside, had similar thickness in mice treated with CR5 and exposed to light as in dark-adapted mice, while in vehicle-treated illuminated mice this layer was ~50-70% shorter (**Fig. 5D**). The retina injury with bright light induces a stress response that stimulates the activation of resident immune cells, which trigger the migration of immune cells from the peripheral circulation into the retina to clear dying photoreceptors. The increase of autofluorescent (AF) spots related to the activation of these immune cells can be detected by an *in vivo* whole fundus scanning laser ophthalmoscopy (SLO) imaging (**Fig. 5E**). The number of AF spots largely increased in the vehicle-treated light-exposed mice as compared to mice kept in the dark. However, the treatment with CR5 resulted in a decrease of AF in the retina (**Fig. 5F**). The retinal function was also preserved in mice treated with a single dose of CR5 prior to illumination. The ERG responses were diminished in vehicle-treated and light-exposed mice as compared to mice kept in the dark. However, both a- and b-wave responses detected in mice administered with CR5 closely resembled the responses detected in unexposed mice (**Fig. 5G**). These results demonstrate that CR5 exhibits a protective effect against light-induced retinal degeneration in *Abca4^-/-^Rdh8^-/-^* mice, possibly due to its opsin-stabilizing property.

**Figure 5.**
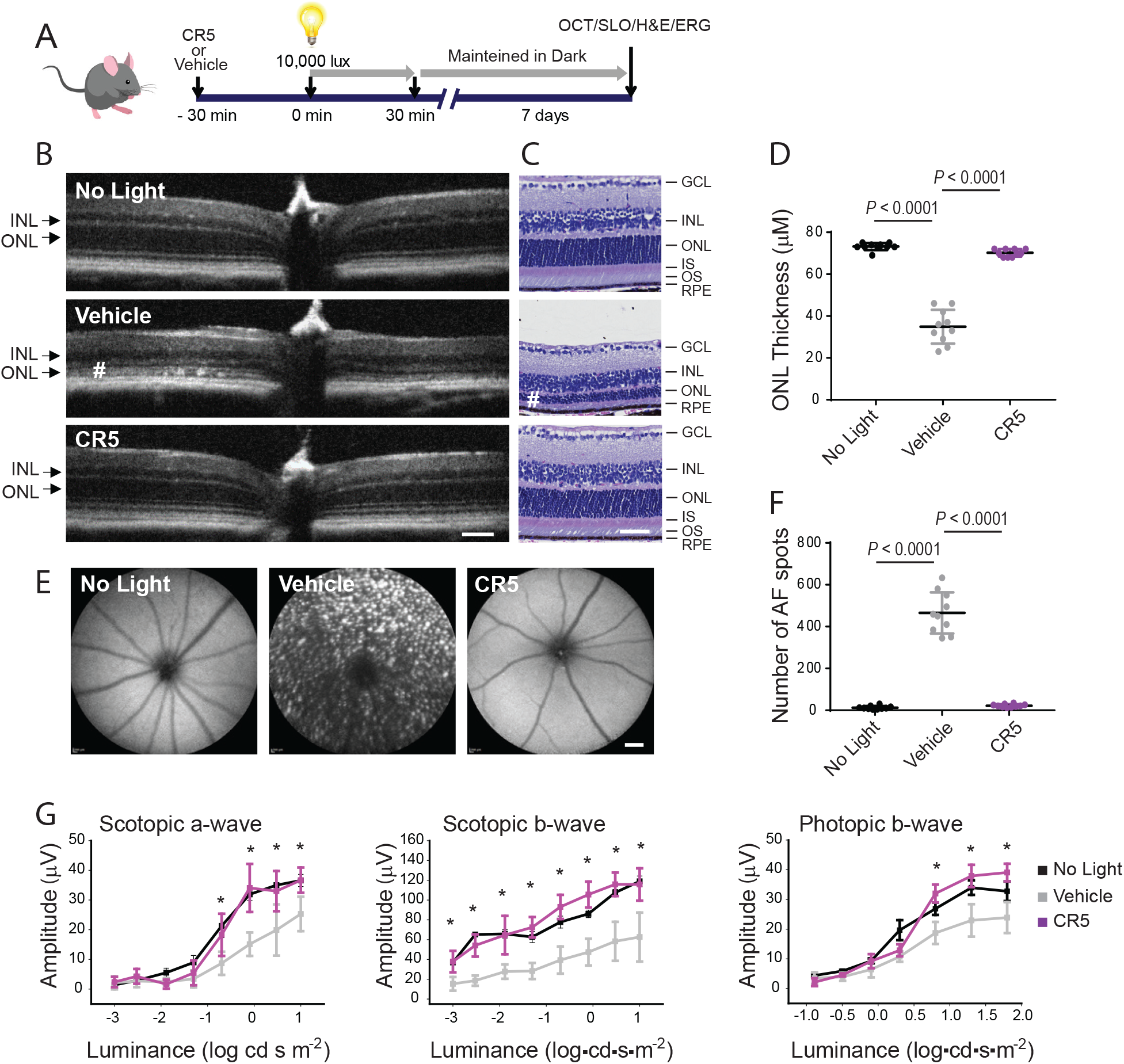
Protective effect of CR5 against light-induced retinal degeneration. **A**, Experimental design. The effect of CR5 on retinal degeneration induced by acute light was investigated in *Abca4^-/-^Rdh8^-/-^* mice vulnerable to bright exposure according to the experimental design described in Material and Methods. **B**, The representative optical coherence tomography (OCT) images of the retina. Shorten photoreceptor cells layer is indicated with (#). Scale bar 100 μm. **C**, The retinal morphology was examined in the hematoxylin and eosin (H&E)-stained paraffin eye sections. Shorten photoreceptor cell layer is indicated with (#). Scale bar 50 μm. **D**, The measurement of the outer nuclear layer (ONL) thickness at 500 μm from the optic nerve head. The measurement was performed in n = 6 mice per treatment group. Statistically different changes were observed in vehicle-treated mice as compared to dark-adapted mice. No statistical difference was found for dark-adapted and CR5-treated mice. Error bars indicate standard deviation (S.D.). *P* values for the statistically different changes (*P* < 0.05) are indicated in the figure. **E**, Representative scanning laser ophthalmoscopy (SLO) images of the whole fundus. Autofluorescence (AF) spots were detected in vehicle-treated mice Scale bar 1 mm. **F**, Quantification of the AF spots was performed in n = 6 mice per treatment group. Statistically different changes were observed in vehicle-treated mice compared to dark-adapted mice and in the CR5-treated mice compared to vehicle-treated mice. No statistical difference was found for dark-adapted and CR5-treated mice. Error bars indicate standard deviation (S.D). *P* values for the statistically different changes (*P* < 0.05) are indicated in the figure. **G**, Retinal function examined by the ERG responses. ERG measurements were performed in n = 5 mice per treatment group. CR5 administered to mice prior to light exposure protected both a-wave and b-wave responses as compared to vehicle-treated mice. These changes were statistically different. No statistical difference was found for dark-adapted and CR5-treated mice. Error bars indicate S.D. Statistically different changes (*P* < 0.05) in the ERG responses after treatment with CR5 compared to vehicle-treated mice are indicated with asterisks. Statistical analyses were performed with the two-way ANOVA and post hoc Turkey’s tests. IS, inner segments; OS, outer segments; INL, inner nuclear layer; GCL, ganglion cell layer; RPE, retinyl pigment epithelium.

We next sought to determine whether CR5 could slow the progression of photoreceptor cell death in RP. The P23H Rho mutation is one of the most prevalent causes of RP worldwide, and heterozygous *knock-in Rho^P23H/+^* mice closely resemble retina degeneration in humans carrying this mutation (34). Moreover, we recently showed that photoreceptor degeneration could be decelerated in these mice upon treatment with flavonoids (19). Thus, we applied a similar administration regimen of CR5 to *Rho^P23H/+^* mice and evaluated its effect on the retinal morphology and function. *Rho^P23H/+^* mice were injected i.p. with CR5 at 10 mg/kg b.w. every other day starting at postnatal (P) day P21 and evaluated at P33 (**Fig. 6A**). The CR5 compound did not cause apparent toxicity as evidenced by the normal body weight growth of these mice during the treatment period in comparison to vehicle-treated mice (**Fig. 6B**). No obvious behavioral changes were also detected. Morphological evaluation of the retinas by using OCT *in vivo* imaging and histological assessment revealed the increased thickness of the ONL in mice treated with CR5 as compared to vehicle-treated mice (**Fig. 6C-F**). The immunohistochemical staining of these tissues suggests the expression of both Rho and cone opsins was increased in mice treated with CR5 as compared to vehicle-treated mice (**Fig. 6E**). Consistent with the improvements of the retina structure, retina function was also noticeably improved in mice treated with CR5 as compared to vehicle-treated mice (**Fig. 6G**). The amplitude of both a- and b-wave scotopic responses and b-wave photopic responses were enhanced in mice treated with CR5. These findings demonstrate that CR5 can prolong the survival of photoreceptors in mice modeling human RP.

**Figure 6.**
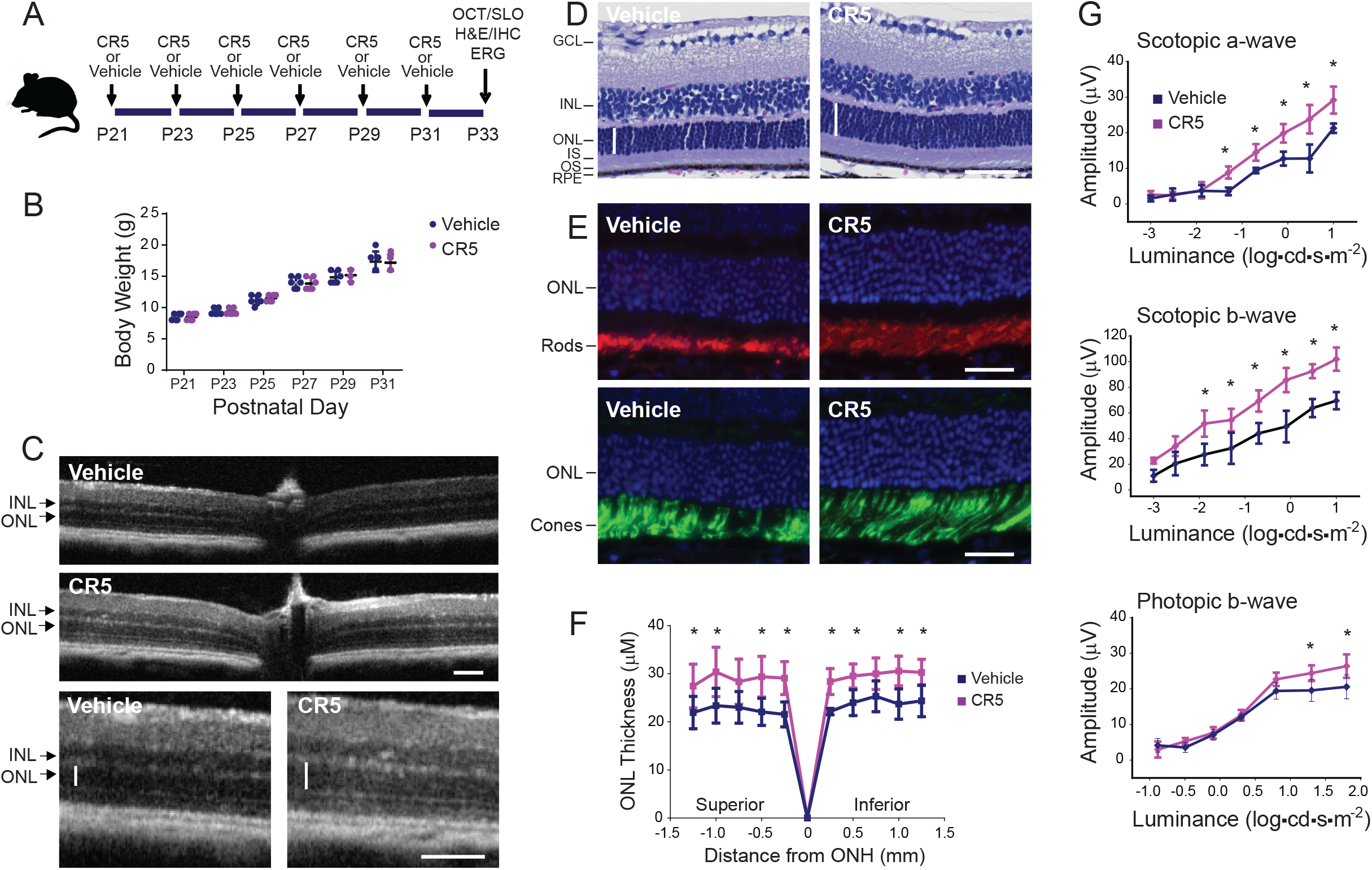
Protective effect of CR5 against the progression of retinal degeneration in RP. **A**, Experimental design. The CR5 compound (10 mg/kg b.w.) or vehicle were administered to heterozygous *Rho^P23H/+^* mice at post-natal day P21 every other day. Mice were analyzed at P33. **B**, Body weight (b.w.) analysis of CR5 or vehicle treated mice. B.w. was measured every other day between P21 and P33. **C**, The retinal morphology of the mouse eyes was analyzed by the optical coherence tomography (OCT) imaging. The representative images are shown. The measurement was performed in n = 6 mice per treatment group. Scale bar, 100 μm. **D**, Images of the hematoxylin and eosin (H&E)-stained retinal sections. Scale bar, 50 μm. **E**, Immunohistochemistry in the cryo-sections prepared from eyes collected from CR5 or vehicle treated mice. Sections were stained with anti-Rho specific antibody (red), showing organization of rod photoreceptors, peanut agglutinin (PNA) (green), showing health of cone photoreceptors, and DAPI, which labels cell nuclei (blue). Scale bar, 25 μm. **F**, The thickness of the retinal outer nuclear layer (ONL) was measured at 0.25, 0.5, 0.75, 1.0, and 1.5 mm from the optic nerve head (ONH). Error bars indicate standard deviation (S.D). The changes found in mice treated with CR5 were statistically different as compared to vehicle-treated mice. The statistically different changes (*P* < 0.05) are indicated with asterisks. **G**, Retinal function was examined by measuring the ERG responses. The ERG measurements were carried out in n = 5 mice per treatment group. Statistically different changes (*P* < 0.05) in the ERG responses after treatment with CR5 compared to vehicle-treated mice are indicated with asterisks. Statistical analyses were performed with the two-way ANOVA and post hoc Turkey’s tests. IS, inner segments; OS, outer segments; INL, inner nuclear layer; GCL, ganglion cell layer; RPE, retinyl pigment epithelium.

## DISCUSSION

Rho is the main light-sensing receptor in the retina. The correct folding of Rho and its transport to the rod outer segment is critical for proper morphogenesis and function of photoreceptors (51,52). The *RHO* gene is prone to spontaneous mutations resulting in the conformationally unstable receptor that is retained in the secretory pathway (2,53,54). Failure to express correctly folded Rho diminishes the supply of the functional receptor, leading at first to night blindness and eventually to blindness in RP (2,3). Most of these mutations destabilize the core interactions involved in the binding of the natural chromophore 11-*cis*-retinal important for Rho stability and correct folding (5,54,55). This is often associated with prolonged Rho regeneration delaying the formation of the functional receptor (56). Treatment of misfolded RP variants with 11-*cis*-retinal or its analogs in the heterologous expression systems promotes the correct routing of mutant receptors to the plasma membrane (5,6). However, the toxicity of retinals limits their use as pharmacological chaperones *in vivo*.

Recent discoveries have indicated that the rod opsin’s orthosteric site could accommodate a variety of other small molecules that could potentially help to re-stabilize misfolded variants (16–18,57). Indeed, we recently found that polyphenolic compounds such as flavonoids bind to rod opsin and enhance its stability (18). For instance, quercetin restores multiple native interactions within the chromophore-binding site of the P23H rod opsin and improves the function of this mutant *in vitro* (19). Moreover, treatment with quercetin prevents retinal degeneration caused by excessive accumulation of unliganded opsin upon exposure to bright light and importantly slows the progression of photoreceptor cell death in the P23H Rho *knock-in* mouse model of RP (19,20). In this study, we followed up on our prior discoveries by carrying out a virtual screening to identify other chemical chaperones with natural product scaffolds that form favorable interactions within the retinal-binding pocket. Our results identified several novel molecules with binding free energies lower than quercetin and predicted drug-likeness that could occupy the orthosteric site. Interestingly, all these molecules contained a common chromenone motif. As shown before, chromenone-based scaffolds display a drug-like structure with favorable physicochemical properties, and offer various possibilities for chemical substitutions in efforts to improve their biological activity (25,58). Compounds featuring a chromenone core structure were found to have agonistic or antagonistic activities for orphan GPCR35 and lipid-activated GPCR55 (25–27). Moreover, the antagonist of cysteinyl leukotriene 1 (CysLT1) receptor marketed as the antiasthmatic drug, pranlukest contains a chromenone moiety in its structure (59,60). The chromenone derivatives identified *in silico* in this current study could bind to rod opsin, modulate its conformation, and increase its stability. However, CR4 and CR5 displayed distinct effects on pigment regeneration with a photostable analog of the native 11-*cis*-retinal ligand. While preincubation of opsin with CR5 reduced binding of 9-*cis*-retinal, CR4 slightly increased retinal binding, which highlights differences in the manner in which these compounds interact with rod opsin. However, both compounds affected the receptor function. Based on the effect on pigment regeneration it was expected that CR5 could modulate Rho function by competing with the retinal ligand for the occupation of the orthosteric binding site.

Encouragingly, these compounds enhanced the surface immunostaining of the P23H rod opsin variant, suggesting that they possess chaperone properties to assist the folding and transport of this misfolding variant from the ER to the plasma membrane. CR5 had a greater effect than CR4, which may reflect different modes of interaction of these compounds within the receptor. An analysis of the effects of these compounds on a wider array of RP mutants by deep mutational scanning indicated that only CR5 measurably increased the plasma membrane expression of 31 mutants that were also stabilized by 9-*cis*-retinal in the control experiment. Interestingly, the mutated residues of the variants rescued by the CR5 compound are involved in the stabilization of the chromophore within the binding pocket (54). The small molecules capable to restore native interactions within the protein core can restore the folding and membrane targeting of pathogenic rod opsin (19,47,55). This result suggests that CR5 possesses chaperone properties recognizing the orthosteric binding site and enabling the shift of the mutant receptor towards a state targetable to the cell surface in the absence of the retinal chromophore. Altogether, the data presented in this study could serve as a foundation to identify the CR5 pharmacophore and based on that propose further chemical modifications to develop new analogs of CR5 with improved binding and chaperone activities for misfolding rod opsin variants. Emerging observations suggest that the potency of pharmacochaperones is typically proportional to their binding affinity (61,62). Thus, future CR5 analogs that bind to rod opsin with higher affinity could potentially offer greater chaperone potency.

Importantly, CR5 showed therapeutic benefit *in vivo,* in two mouse models of retinal degeneration. *Abca4^-/-^Rdh8^-/-^* mice are susceptible to light-induced retinal degeneration due to a prolonged presence of ligand-free rod opsin and delayed clearance of all-*trans*-retinal photoproducts (32). As we showed, the stabilization of opsin with either 11-*cis*-6-member lock retinal analog or nonretinoid small molecule stabilizers protects photoreceptors in these mice exposed to bright light insult (16,35). Interestingly, CR5 administered before light illumination showed a similar protective effect for retina health in these mice. Moreover, CR5 prolonged the survival of photoreceptors in RP-related *knock-in* P23H mice, likely due to its ability to increase the conformational stability of mutated rod opsin, which assisted the correct folding and transport of the mutant receptor. As we recently demonstrated, quercetin provided similar advantageous effects slowing the progression of RP in these mice (19).

In conclusion, in this study, we have identified novel opsin stabilizing chromenone-containing small molecules with chaperone properties. Based on our cumulative observations, CR5 could be a valuable lead compound with therapeutic potential for retinal dystrophies associated with photoreceptors degeneration.

## ACKNOWLEDGEMENTS

This work was supported by funding from the National Institutes of Health EY025214 (BJ) and R01GM129261 (JPS). The authors thank the Visual Science Research Center Core at Case Western Reserve University (supported by the NIH grant P30 EY011373) with special gratitude directed to Catherine Doller for assistance with tissue sectioning and H&E staining and Maryanne Pendergast for assistance in slide imaging.

## AUTHOR CONTRIBUTIONS

B.J. and J.T.O. conceived and designed the experiments.

J.T.O, B.J., A.G.M, F.J.R, W.D.P., and J.P.S. conducted the experiments.

B.J., J.T.O., A.G.M, F.J.R, W.D.P., and J.P.S. contributed to the writing of the manuscript.

B.J. and J.P.S coordinated and oversaw the research project.

All authors discussed the results and commented on the manuscript.

## ABBREVIATIONS

adRP: autosomal dominant retinitis pigmentosa
BBB: blood-brain barrier
BRB: blood-retina barrier
bw(s): body weight(s)
DAPI: 4’6’-diamidino-2-phenyl-indole
CSNB: Congental Stationary Night Blindness
DMEM: Dulbecco’s modified Eagle’s medium
DMSO: dimethyl sulfoxide
EDTA: ethylenediaminetetraacetic acid
FBS: fetal bovine serum
GPCR: G protein-coupled receptor
PBS: phosphate-buffered saline
PNA: peanut agglutinin
PVDF: polyvinylidene difluoride
ROS: rod outer segments
RP: retinitis pigmentosa
RT: room temperature
S.D.: standard deviation
WT: wild type.

