## Supplemental data for "Chromenone derivatives as novel pharmacological chaperones for retinitis pigmentosa-linked rod opsin mutants"

**RUNNING TITLE:** Novel modulators of rhodopsin mutants

[#] **CORRESPONDENCE**

Beata Jastrzebska, Ph.D., Department of Pharmacology, School of Medicine, Case Western Reserve University, 10900 Euclid Ave., Cleveland, OH 44106-4965, USA; Phone: 216-368-5683; Fax: 216-368-1300;, ORCID ID: https://orcid.org/0000-0001-5209-8685.

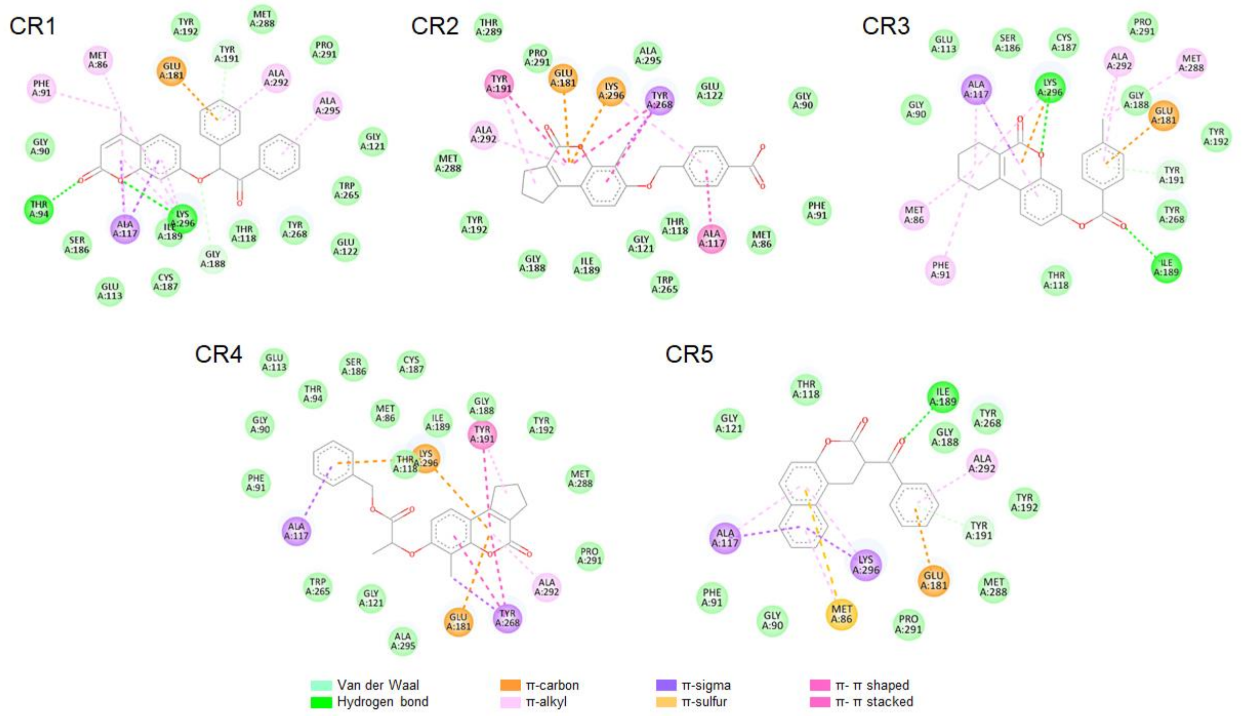


**Figure S1. Two-dimensional representation of the interaction between the new compounds and the residues in the opsin's polypeptide chain.** These are 2D representations of the low energy structures obtained from molecular docking of CR1-CR5 compounds to the rod opsin orthosteric site and visualized with the Biovia Discovery Studio Visualizer 17.2.0 visualizer.

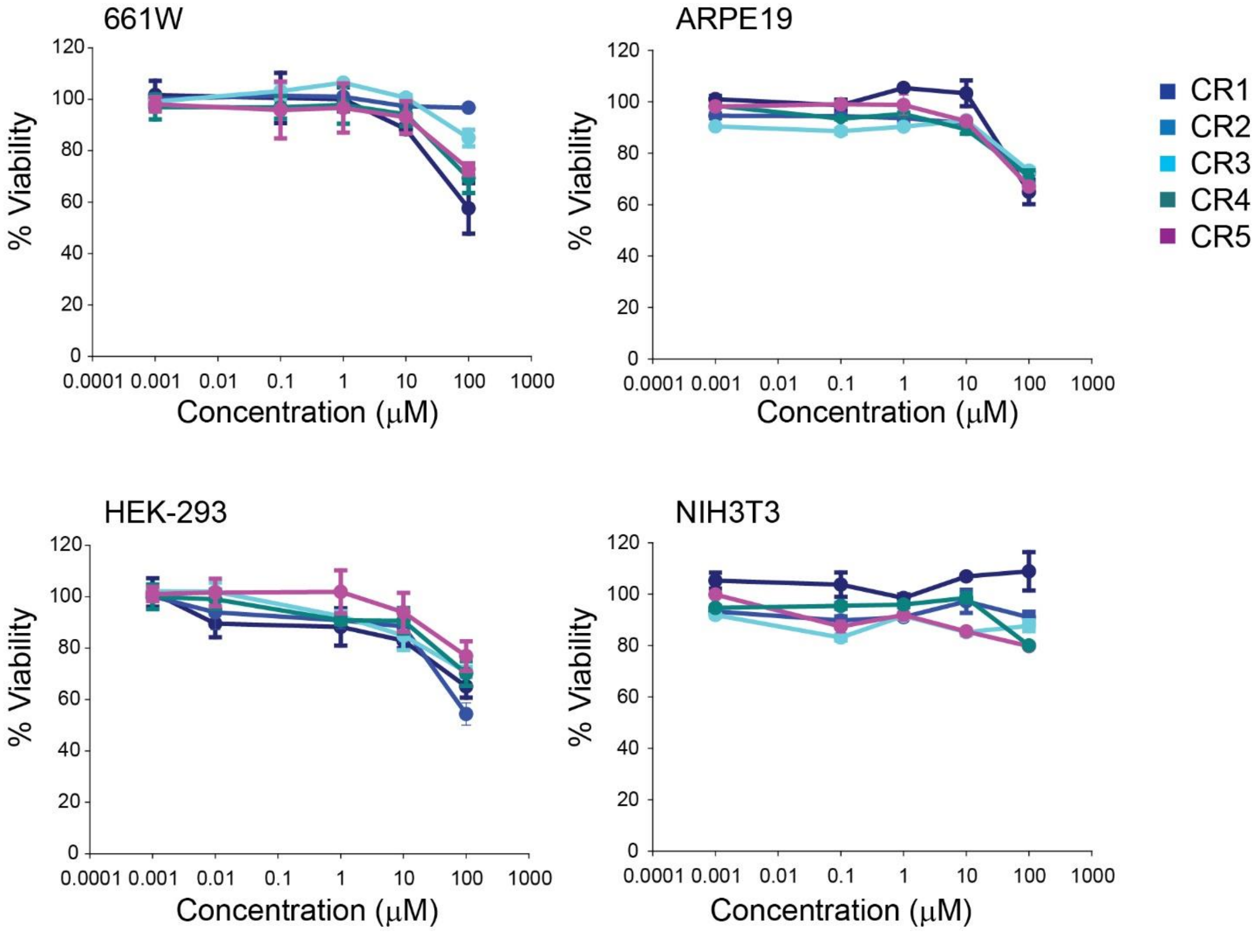


**Figure S2. Cytotoxicity of the new compounds.** Toxicity of CR1-CR5 compounds was tested in four cell lines, the NIH-3T3, HEK-293 cells, photoreceptor-derived 661W cells, and retinyl pigment epithelial (RPE)-derived ARPE19 cells by using MTT assay treated with the compounds at 0.001-100 μM concentration for 24 h. The results are presented as a percentage of viable cells as compared to non-treated control cells.

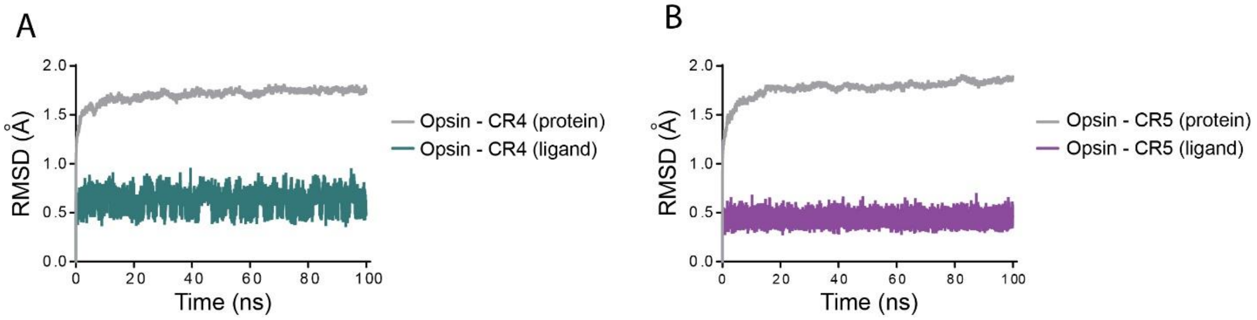


**Figure S3. Stability of CR4 and CR5 binding within the orthosteric site of rod opsin.** A monomer of wild type (WT) rod opsin (PDB ID: 3CAP) was used to perform molecular dynamics simulations (MD) to assess the binding stability of the CR4 (**A**) and CR5 (**B**) within the orthosteric site of rod opsin and their effect on receptor conformational stability.
